# FASER: A TOOL TO SIMULATE PSF DISTORTIONS IN STED MICROSCOPY

**DOI:** 10.1101/2024.09.27.615327

**Authors:** Stephane Bancelin, Johannes Roos, U. Valentin Nägerl

## Abstract

We introduce Faser, a software package developed in Python as a plugin for the open-source napari platform, designed to simulate the excitation point spread functions (PSFs) of microscopes. Using a full-vectorial computational approach to simulate the electromagnetic fields within the focal region, it makes precise predictions and allows detailed analyses of excitation PSFs. Faser is intended as a pedagogical tool enabling users to explore the impact of various geometrical and optical parameters of practical importance on imaging performance. It supports the modeling of complex beam profiles, including donut and bottle-shaped beams, which are instrumental in advanced microscopy techniques such as Stimulated Emission Depletion (STED) microscopy. Through specific simulations and accessible illustrations, we showcase Faser’s capabilities in replicating the distinctive properties of STED beams, making it a valuable resource for researchers and students in optical microscopy to explore and optimize high-resolution imaging techniques.

## 1 Introduction

Optical microscopy occupies a central place in the field of biophotonics, with new imaging technologies enabling significant biological breakthroughs. As technology keeps progressing, new variations and technical improvements are being continually developed, pushing the limits of optical microscopy in terms of spatial and temporal resolution [1], sensitivity, specificity, non-invasiveness and depth penetration [2], heralding the era of quantitative multiscale optical biology [3, 4].

Optical microscopy typically relies on the use of tightly focused light fields. In most cases image quality (*e*.*g*. SNR, spatial resolution, sensitivity) directly relates to the spatial shaping of the Point Spread Function (PSF) of the microscope. Hence, precise determination of the optical properties of focused beams (intensity, polarization and phase) is critical for designing optical microscopes and achieving high imaging performance. Beyond the optical beam properties, the PSF can be strongly affected by any heterogeneity in the optical path, whether in the microscope or the sample) [5, 6, 7].

In this context, the study of how to focus light in the best way has drawn considerable interest for many decades. Over the years, an extensive literature on analytical and numerical models to compute the diffracted light distribution has emerged. Since high resolution optical microscopy and nanoscopy commonly employ high numerical aperture (NA) objectives, a fully vectorial model is required to described the focused light field. The corresponding equations have been well established for years [8, 9], notably through the work of P. Török [10, 11], with analytical or numerical solutions investigated in specific configurations [12, 13, 14].

However, a general analytical solution has not yet been derived. Due to the complexity of numerical calculations, various approximations are often used, limiting the simulation of the PSF to specific cases. In particular, paraxial approximation (small angles with respect to the optical axis), scalar approximation (neglected polarization of light), and aberration-free conditions are commonly considered.

Here we introduce Faser, an open-source Python software package designed to simulate the PSF of an optical microscope using fully vectorial calculations. In particular, it allows users to set the intensity profile, the polarization and a phase mask of the incident laser beam, as well as modify the geometry of the focusing medium. This tool has been designed for pedagogical purposes, enabling users to simulate the effect of key optical and geometrical parameters on the PSF of the microscope. Specifically, Faser is well-suited for *in vivo* experiments, making it possible to simulate the impact of a cranial window (*e*.*g*. opening, coverslip, tilt). This is instrumental for identifying and defining the critical parameters for designing and optimizing an optical imaging protocol.

Developped as a Plugin for Napari [15], Faser provides a graphical user interface (GUI) that allows running simulations directly without any coding knowledge. Additionally, a batch modes functionality, defined for most parameters, enables users to vary parameters systematically to determine their impact and tolerance range.

In this work, we illustrate the potential of Faser to simulate shaped PSFs, such as those used in STED and RESOLFT microscopy, in the context of *in vivo* microscopy, including a cranial window and a coverslip. We investigate the effects of common cases of misalignments and optical aberrations on the STED PSF and on the resulting effective fluorescence PSF.

## 2 Theoretical background

### 2.1 Light distribution in the focal region

In an optical imaging system, if the NA of the focusing lens is relatively high (NA ≥ 0.7), the approximations used in scalar diffraction theory, such as the paraxial approximation, Kirchhoff boundary condition and Fresnel or Fraunhofer approximation are no longer valid [8]. In the 1950’s, Richard and Wolf proposed a comprehensive mathematical representation of the electromagnetic field (EM) distribution in the focal region of a high NA objective lens [16, 17]. The main assumptions of this theory can be summarized as follows:

- The beam at the exit pupil has a spherical wavefront with a radius *f*, the focal length of the objective,
- Each diffracted light ray is considered as a plane wave propagating towards the geometrical focal point of the lens, indicated by the wave vector **k**.
- The angle *β* between the optical axis and the propagation direction of the diffracted ray is small (cos(*θ*) ≈ 1).

Following this formalism, known as the vectorial diffraction theory, the electric field (**E**) in an arbitrary point P(*x, y, z*) in the focal region of a high NA objective lens results from the superposition of all diffracted plane waves emerging from the exit pupil of the lens within the solid angle Ω [17, 8, 10]:

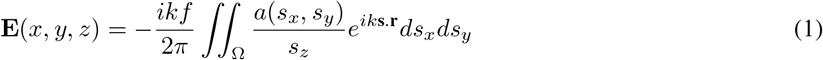

where k is the wavenumber, **s** = (*s*_*x*_, *s*_*y*_, *s*_*z*_) is a dimensionless unit vector along each ray from the objective pupil towards the geometrical focal point O, *a* the complex amplitude of the incident laser beam after the objective and **r** the position vector of point P(*x, y, z*). For the sake of clarity, it is more convenient to express the wave vector **s** and pupil function in spherical coordinates. In this case:

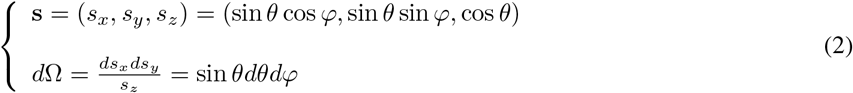

Considering the geometry depicted in Fig. 1, the diffraction integral can be expressed, as :

**Figure 1:**
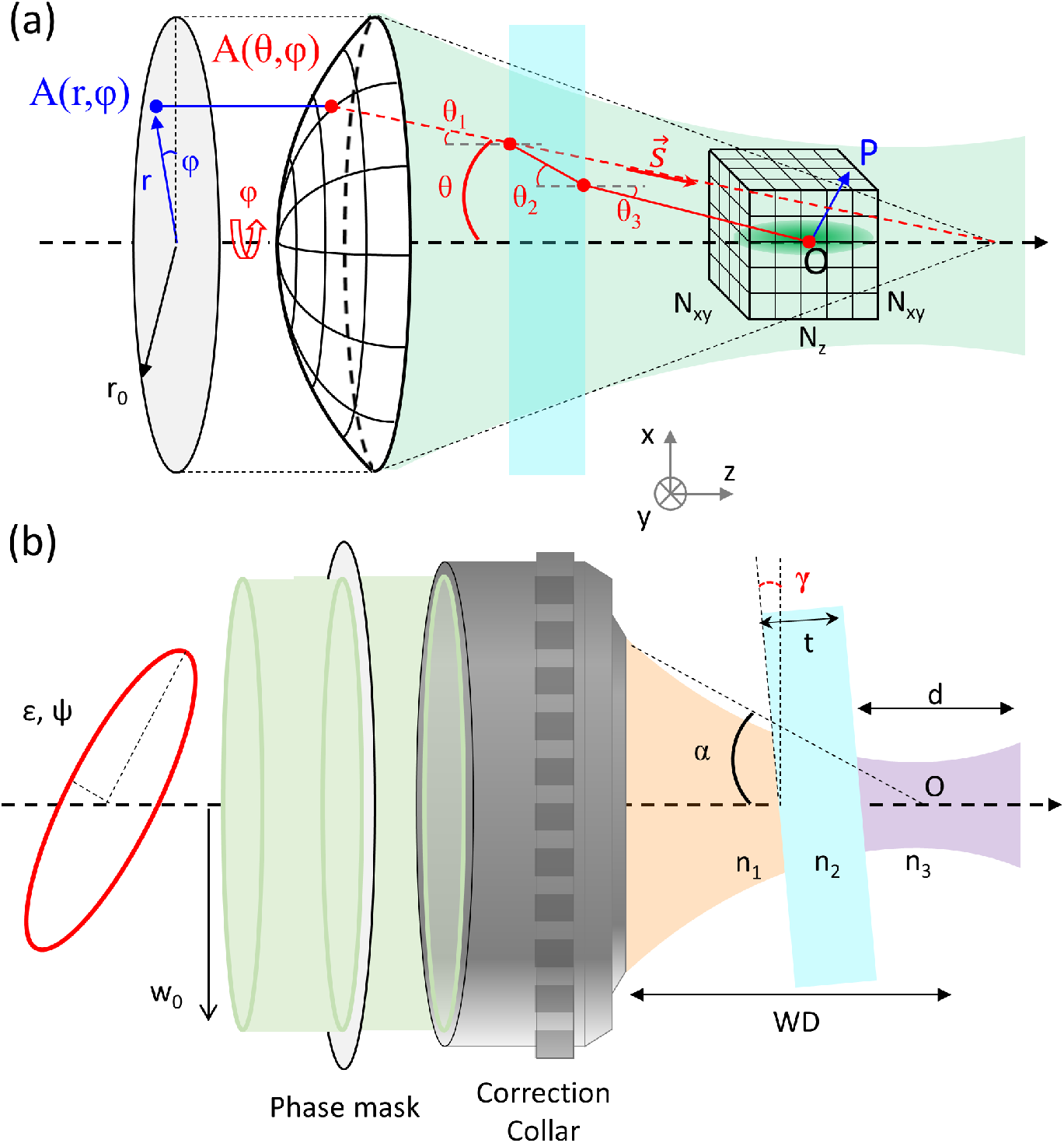
Schematic of the geometry used for the simulations. a) Using vectorial diffraction theory, the incident pupil field *A*(*r, φ*) is transformed into a spherical wavefront through the objective lens and then propagated to the focal spot. The focal intensity is calculated in any point *P* in the vicinity of the focus *O*. The focalization process is computed through the stratified media consisting of the immersion liquid, the coverslip and the sample. b) The ellipticity, intensity profile and potentials aberrations in the input beams are determined before the objective. A phase mask can also be added in the context of beam shaping, notably STED microscopy.

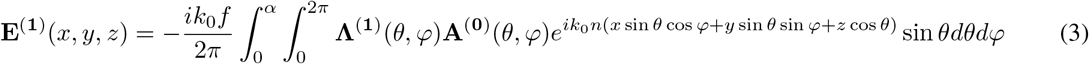

with *k*_0_ the wavenumber in the vacuum, n the refractive index in the medium, *α* = arcsin(NA*/n*) the marginal ray angle of the objective (semi-aperture angle) and where **Λ**^(**1**)^(*θ, φ*) is an operator describing the transformation of the complex vector field **A**^(**0**)^(*θ, φ*) incident at the back aperture of the objective lens into the field **A**^(**1**)^(*θ, φ*) on Ω after refraction by the objective lens.

### 2.2 Focusing through a glass coverslip

Experimentally, in optical microscopy, light is typically focused through water or oil immersion media and a coverslip is commonly used to protect the sample. To investigate this configuration, Riachard and Wolf’s solution has been later extended by Törok *et al*. [10] to account for a planar dielectric interface and further generalized to stratified media [11]. Although this framework can describe an arbitrary number of layers with different refractive indices, only two interfaces are necessary to describe a glass coverslip placed between the sample and the immersion media Fig. 1.

In this case, the focal field in the sample, after the coverslip can be written:

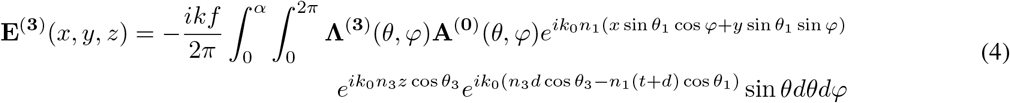

with *d* the depth in the sample, *t* the thickness of the coverslip and *n*_*l*_ and *θ*_*l*_ the refractive indices and incident angles in the immersion medium (1), coverslip (2) and sample (3), respectively. These parameters are determined according to Snell’s law.

Note that **Λ**^(**3**)^(*θ, φ*) describes the polarization conversion in an heterogeneous medium after the objective aperture, including Fresnel coefficients of reflection and transmission through the coverslip. Notably, it includes the different apodization for *s* and *p* polarization.

The third exponential term corresponds to aberrations induced along the optical path through the coverslip and the biological sample. Notably, it is independent of the polar angle *θ*_1_ and thus describes spherical aberrations. The effect of a correction collar can hence be simulated, assuming perfect correction mechanisms, by applying the inverted aberration terms:

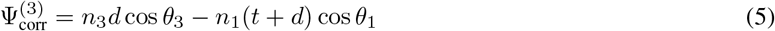

## 3 Faser: vectorial PSF simulations

As highlighted in Eq. 4, the PSF in the focal region of a high NA objective lens depends on several parameters. Although these equations have been well-known for decades, performing full vectorial simulations of the PSF in the focus of a high NA objective lens remains a complex task, and up to now, most freely available simulation tools only allow calculations for specific configurations.

Faser is intended as a pedagogical tool for optical afficionados, making it possible to explore the impact of numerous parameters related to the excitation beam and the focusing geometry. This flexibility facilitates the simulations of PSFs that closely reflect real-world experimental conditions. The following subsections outline these parameters and how they are implemented in Faser.

### 3.1 Numerical calculation

In principle, there is an infinite number of diffracted rays within the solid angle Ω that emerge from the lens aperture and propagate towards the focal region. Yet, for numerical computations, the polar *θ* and azimuthal angles *φ* are discretized into *N*_*θ*_ and *N*_*φ*_ steps, equally spaced by Δ*θ* and Δ*φ* respectively:

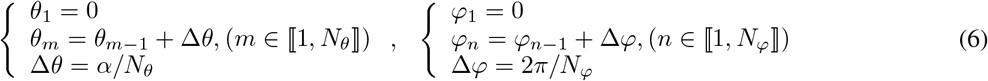

Consequently, the diffraction integral may be discretized by summing up a finite number of plane waves. This discretized form is more convenient for numerical computation and can be written as:

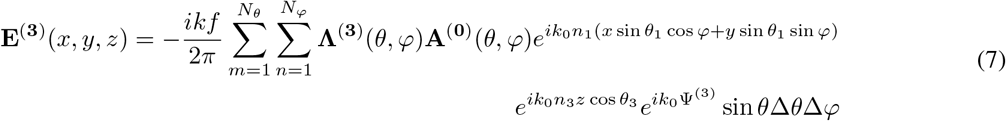

The focal intensity is then calculated as the squared modulus of the electric field:

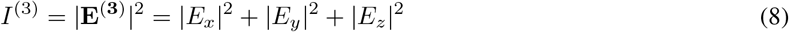

### 3.2 Incident beam

The incident beam on the input pupil of the objective lens can be written :

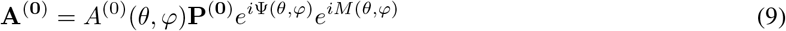

where *A*^(0)^(*θ, φ*) is the complex amplitude, **P**_**0**_ the polarizatioon of the incident beam, Ψ(*θ, φ*) the aberration function and *M* (*θ, φ*) the phase profile of the input beam, corresponding to the phase mask used to shape the beam.

#### 3.2.1 Amplitude

The amplitude of a Gaussian beam incident on the input plane of the objective lens can be written in cylindrical coordinates as:

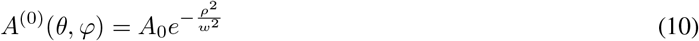

where *A*_0_ is a constant and *w* is the beam waist. While passing through an aplanatic objective, the incident plane wave transforms into a spherical wave converging to the focal point. Therefore, the amplitude distribution after the objective can be expressed as:

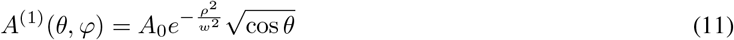

where the cylindrical coordinate on the exit pupil of the objective lens is given by the sine condition [9] *ρ* = *f* sin *θ* and 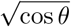 the apodization term ensuring energy conservation while the beam passes through the objective.

### 3.3 Polarization

Assuming a transverse polarization state of the input field, the incident field can be written:\

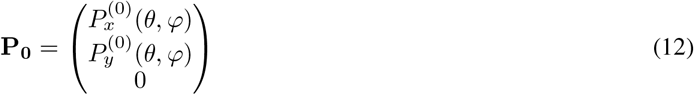

Because of the tight focusing, the objective transforms this input transverse polarization into a partially axial polarization in the focal region polarization, which can be described by the 3×3 lens operator matrix ℒ_*θ*_.

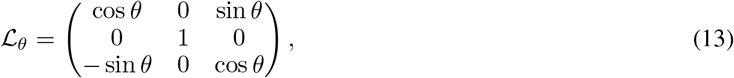

Accounting for the interface of the coverslip, the electric field polarization in the sample (3rd medium) can be written as [11]:

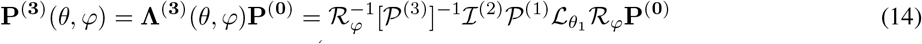

where ℛ_*φ*_ represents the rotation matrix around z axis, 𝒫 (*l*) corresponds to the coordinate system rotation in medium *l* and ℐ ^(2)^ is the matrix describing the effect of the coverslip, considered as a stratified medium of two interfaces.

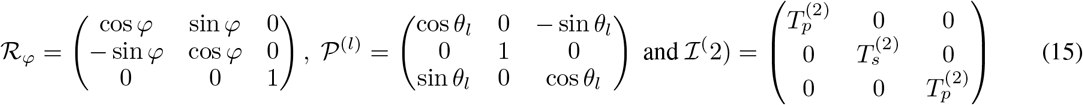

where *θ*_*l*_ is the focusing angle in the *l*^*th*^ medium and 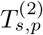 are the transmission coefficients in the coverslip.

In the general case, determining the reflection and transmission coefficients through a stratified medium involves solving Maxwell’s equations. However, in the case of a coverslip with only two interfaces, this simplifies to the well-determined Fresnel coefficients for *-s* and *-p* polarization:

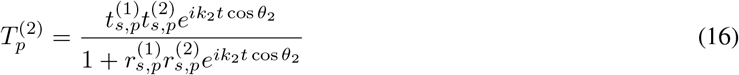

with *t* the thickness of the coverslip and 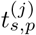 and 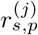 respectively the amplitude transmission and reflection coefficient at the *j*^*th*^ interface:

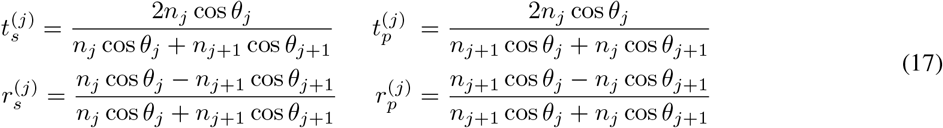

with *n*_*j*_ and *θ*_*j*_ the refractive index and the incident angle in the *j*^*th*^ medium respectively.

For example, in the case of left-handed circular polarization:

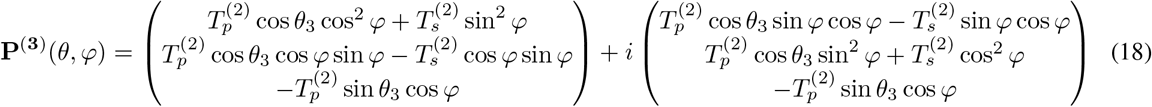

#### 3.3.1 Optical aberrations

In eq. (6), Φ (*θ, φ*) represents the wavefront distortion relative to the reference sphere, an effect commonly called optical aberrations. A convenient approach is to express these aberrations in cylindrical coordinates on the objective pupil and to decompose them using Zernike polynomials:

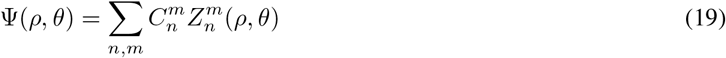

where *n* ∈ ℕ is the radial degree, *m* ∈ ℤ is the azimuthal degree, 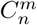 and 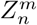 are the Zernike coefficients and polynomials respectively (see table 1).

**Table 1:** First orders Zernike polynomials.

| n | m | Formula | Aberration type |
| --- | --- | --- | --- |
| 0 | 0 | 1 | Piston |
| 1 | 1 | $2\rho \cos(\theta)$ | Tip |
| 1 | -1 | $2\rho \sin(\theta)$ | Tilt |
| 2 | 0 | $\sqrt{3}(2\rho^2 - 1)$ | Defocus |
| 2 | 2 | $\sqrt{6}\rho^2 \cos(2\theta)$ | Astigmatism |
| 2 | -2 | $\sqrt{6}\rho^2 \sin(2\theta)$ | Astigmatism |
| 3 | 1 | $\sqrt{8}(3\rho^3 - 2\rho) \cos(\theta)$ | Coma |
| 3 | -1 | $\sqrt{8}(3\rho^3 - 2\rho) \sin(\theta)$ | Coma |
| 3 | 3 | $\sqrt{8}\rho^3 \cos(3\theta)$ | Trefoil |
| 3 | -3 | $\sqrt{8}\rho^3 \sin(3\theta)$ | Trefoil |
| 4 | 0 | $\sqrt{5}(6\rho^4 - 6\rho^2 + 1)$ | Spherical |

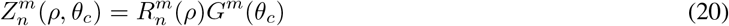

with

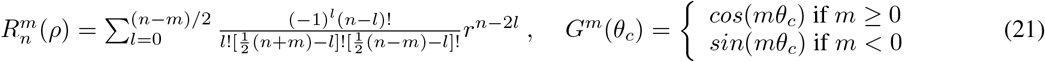

These modes are normalized to 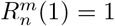 which was introduced by Noll *et al*., in 1975 [18].

Zernike polynomials are widely employed to model optical aberrations due to their orthogonality (on the unit sphere). In the paraxial approximation, each polynomial function corresponds to distinct optical aberration, such as astigmatism, coma, or spherical [19, 20].

Table 1 lists the first order polynomials:

#### 3.3.2 Phase mask

Spatial shaping of the focal field distribution is often achieved using a phase mask *M* (*θ, φ*), imprinted to the incident laser beam prior to reach the objective lens.

Various phase masks are implemented to achieve different spatial shapes. For convenience, the phase mask is expressed in spherical coordinates:

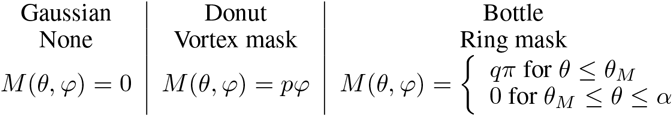

where *q* ∈ ℤ is an integer representing the topological charge of the vortex or ring respectively. To produce a bottle beam *q* has to be odd number. *θ*_*R*_ is the angle between the optical axis and the edge marginal ray of the *π*-phase ring of the phase mask.

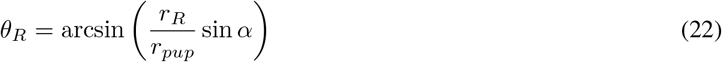

with *r*_*R*_ the radius of the *π*-phase ring and *r*_*pup*_ the radius of the front pupil of the objective.

Finally, the case of donut and bottle beam shaping, which are often used in techniques like 3D-STED microscopy, the two PSFs are treated separately and recombined in the far field using a weighted sum:

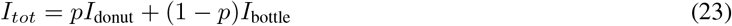

with *p* ∈ [0, 1] so that *p* = 0 and *p* = 1 correspond to a donut- and bottle-shaped beam, respectively.

### 3.4 Effective fluorescence PSF

In the case of STED microscopy, the effective fluorescent spot results from the interaction between the excitation PSF *I*_*e*_ (*eg* confocal, 2-photon) with the de-excitation PSF *I*_*d*_ (*eg* donut, bottle). In Faser, both are computed independently and the effective fluorescent PSF *I*_*eff*_ is determined in every pixel as:

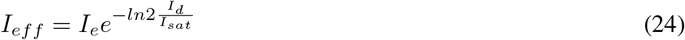

where *I*_*sat*_ represents the saturation factor accounting for the interaction between the STED laser (de-excitation) and the fluorophore.

### 3.5 Focusing geometry

#### 3.5.1 Impact of the cranial window

In many cases, getting optical access to the sample requires placing a coverslip above the region of interest. This practice is prevalent in neuro-imaging, where the use of a ‘cranial window’ enables imaging the brain of living animals. To simulate this experimental configuration, we introduced an additional amplitude mask:

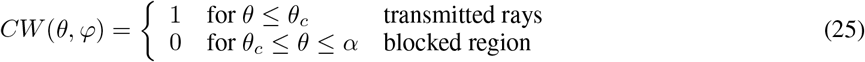

where *θ*_*c*_ is the angle between the optical axis and the marginal ray of the cranial window on the output pupil of the objective. For further details, refer to [7].

In the diffraction integral, this simply leads to a change of the integration limit to accommodate the reduced effective pupil size.

#### 3.5.2 Case of tilted coverslip

In real-world conditions, achieving near-perfect orthogonality of the coverslip with respect to the optical axis can be challenging and in many instance a slight angle, up to few degrees, can easily be present, particularly in the context of *in vivo* imaging. A tilted interface breaks the rotational symmetry of the previous configuration and is therefore expected to affect the phase of the laser beam. Apart from the spherical aberrations induced by the stratified medium from the objective to the sample, a tilted coverslip introduces significant aberrations, notably coma. However, only a few studies deal with this case, most of them using undisclosed algorithms. In Faser, we implemented an approach proposed by Berning [21] and further detailed in [22] to extend the theoretical framework presented in Section 2.2 to this new geometry.

Schematically, tilting the coverslip by an angle *γ* is equivalent to rotating the objective lens by an angle − *γ*. However, analytically it is simpler to derive the focal field using the rotated objective lens method. Indeed, tilting the coverslip requires to modifying Eq. (4), and notably the integration range and the bounding conditions. In contrast, the second method can simply be realized through a transformation of the pupil function to rotate the spherical wavefront *A*^(1)^ around the origin 0. The integration limit on *θ* is expanded to *α* + |*γ*|, to ensure the entire wavefront is accounted for without clipping, while the part of the beam falling out of the pupil is set to 0 amplitude. The new pupil function 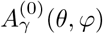 is essentially the pupil function of the untilted case, but evaluated at the transformed position *A*^(0)^(*θ*^*′*^, *φ*^*′*^). The rotated coordinate can be expressed as follow:

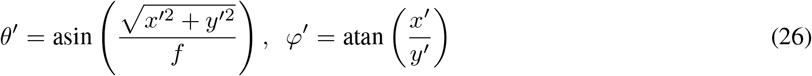

with

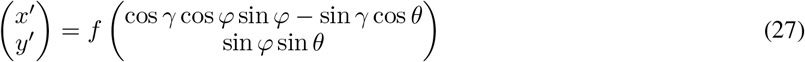

All parameters related to objective lens, and hence to its coordinate system, have to be transformed in this way, notably the apodization factor and the phase correction term Eq. 5.

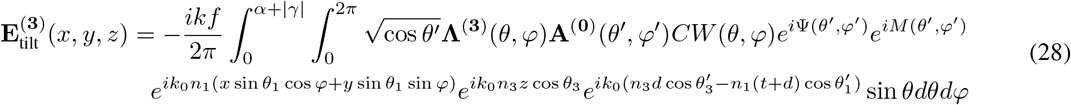

## 4 Installation

Faser provides high-level functions to configure and execute simulations of the focused field. An early design decision of Faser was to package it as a Napari plugin. Faser’s graphical user interface (GUI) is shown in Figure 2. To support the easy integration of Faser into larger automated computational workflows (e.g. when running multiple simultations to explore a larger parameter space), Faser is also executable through a command line interface that comes provided with the “faser” python package.

**Figure 2:**
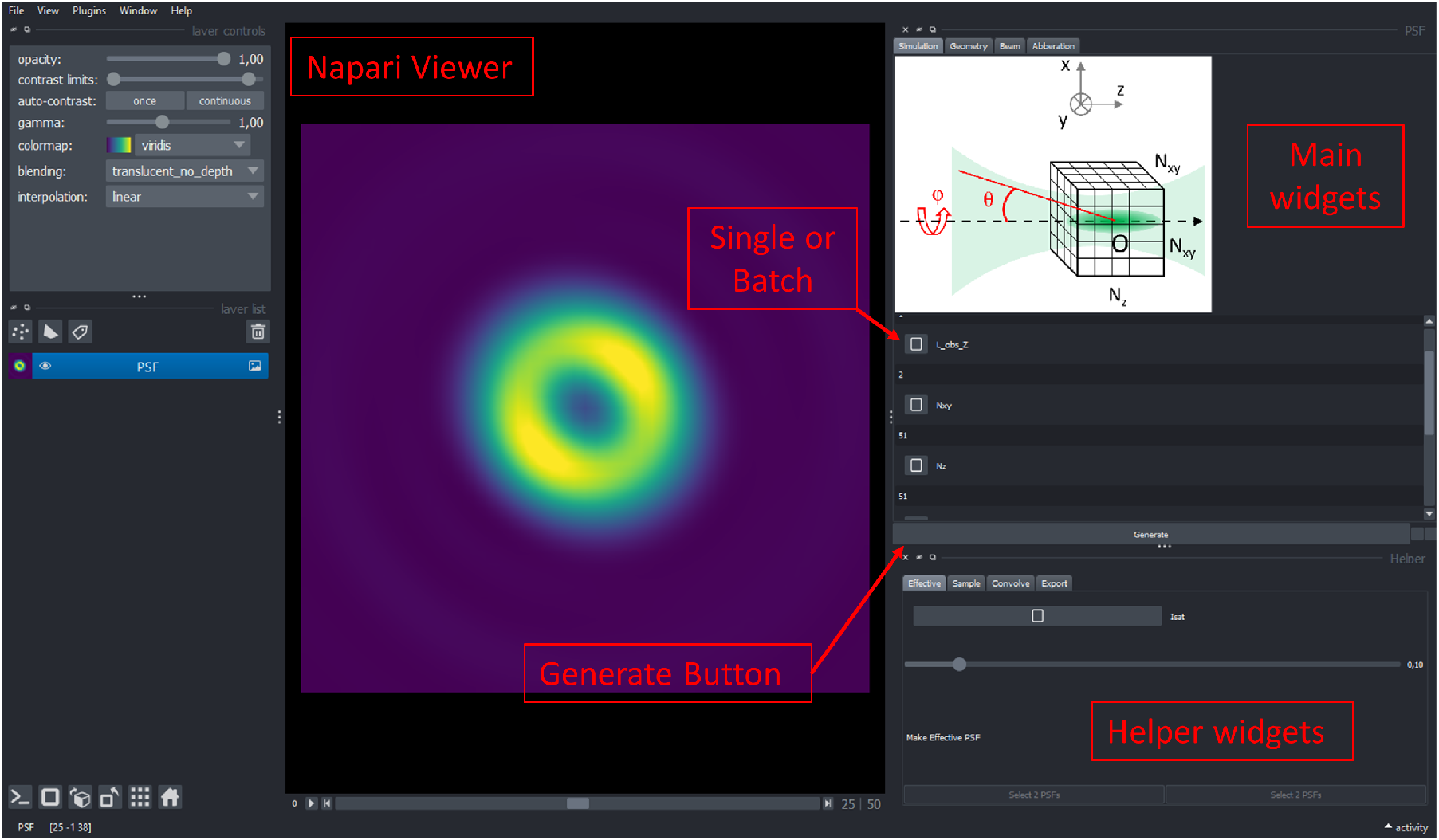
Faser GUI. Based on Napari viewer, Faser features a main widget window with multiple tabs to set the simulation parameters, the property of the input beam and define the focusing geometry. A helper widget window provide additional tabs to compute the effective PSF in STED microscopy, simulate a space and convolve it with the PSF and export the simulated stacks of images.

The source code of Faser is available on https://github.com/jhnnsrs/faser. To install the Faser plugin for Napari from its Github repository, first create a new environment with Python 3.10.11 and pip install Napari. Then, clone the Faser repository by running git clone https://github.com/jhnnsrs/faser.git in your terminal. Navigate to the cloned directory with *cd faser* and install faser using *pip install*, to install the dependencies. Finally, Napari using *python nap*.*py*.

## 5 Faser design and implementation - example for STED microscopy

The objective of Faser is to provide an open-access, educational tool for simulating the excitation PSF of a microscope. The GUI simplifies setting calculation parameters that define the simulation, key parameters including the input beam, the geometry of focalization and the numerical computation parameters.

As an example, in the following sections we focus on Faser’s capability to simulate toroidal PSFs used in STED microscopy. However, as detailed in sections 2 and 3, Faser is based on the vectorial diffraction theory, which offers a general framework for computing the PSF of any optical system. Therefore, the examples presented here could be extended to other microscopy techniques. It is important to note that currently, Faser only considers excitation PSFs. Other parameters, such as pinhole diameter in confocal microscopy, would need to be included to compute the emission PSF.

### 5.1 Main widgets

The main widget contain 4 tabs designed for configuring simulated parameters, input beam properties - including the potential aberrations stemming from flawed optics and setup alignment before the objective lens - and setting the focalization geometry under consideration.

#### 5.1.1 Simulation parameters

The first tab is used to set simulation parameters such as the number of pixels in X, Y and Z directions, along with the observation scale. In addition, the number of integration steps in *φ* and *θ* angles are defined here. A check box also allows normalization of the maximum pixel value of the PSF intensity to 1.

Figure 3 illustrates the effects of these parameters on computation time and PSF accuracy. As expected, computation time increases quadratically with the number of pixels in XY direction. Conversely, changing the number of integration steps *N*_*θ*_ from 5 to 15 strongly impacts the simulated PSF, particularly the quality of the null in the center. Larger number of steps have a more limited impact on the results. However, it is worth noting that the optimal number of integration steps depends on the complexity of the phase mask and the aberration profile of the incident beam. Through various configurations, we found a good compromise between computation time and number of pixels and steps. Yet, simpler PSFs (such a Gaussian beam) can be appropriately computed with fewer steps, whereas highly structured PSFs might necessitate higher numbers in the simulation parameters.

**Figure 3:**
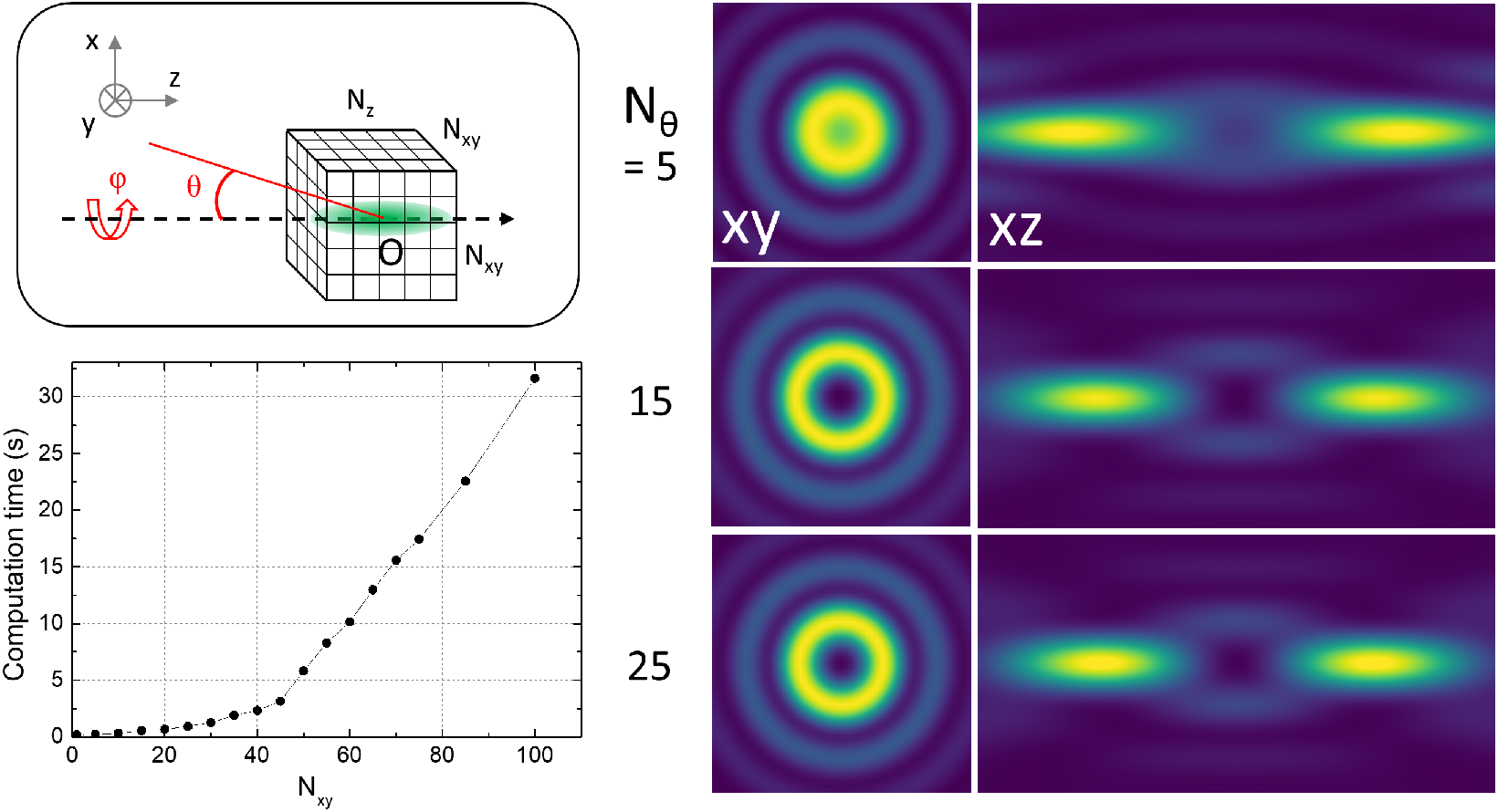
Simulation parameters. a-) Schematic of the simulation configuration. b) Computation time as a function of the number of XY pixels in the computed volume. c) Lateral (2×2 *µm*^2^) and axial (2 × 4 *µm*^2^) views of a *bottle-shaped* PSF for different *θ* integration steps.

To facilitate efficient computation in the Python ecoystem, Faser employs strategies of Just-In-Time Compilation (utilizing Numba) and employs lazy compute patterns through Dask. The computing times provided in the next section has been obtained on a standard desktop running on Windows 11 Pro 64bits, equipped with an Intel core i5-1335U processor (1.30GHz) and 16Gb of RAM memory.

#### 5.1.2 Focusing geometry

The second tab serves to define the focalization geometry under consideration, as depicted in Fig 4. This include specifying the objective parameters (NA, working distance, refractive index of the immersion media and correction collar), details about the potential coverslip (refractive index, thickness, tilt) and the characteristics of the sample (refractive index, depth). In addition, to account for the potential presence of a cranial window used for *in vivo* imaging, an aperture is added right after the objective lens.

**Figure 4:**
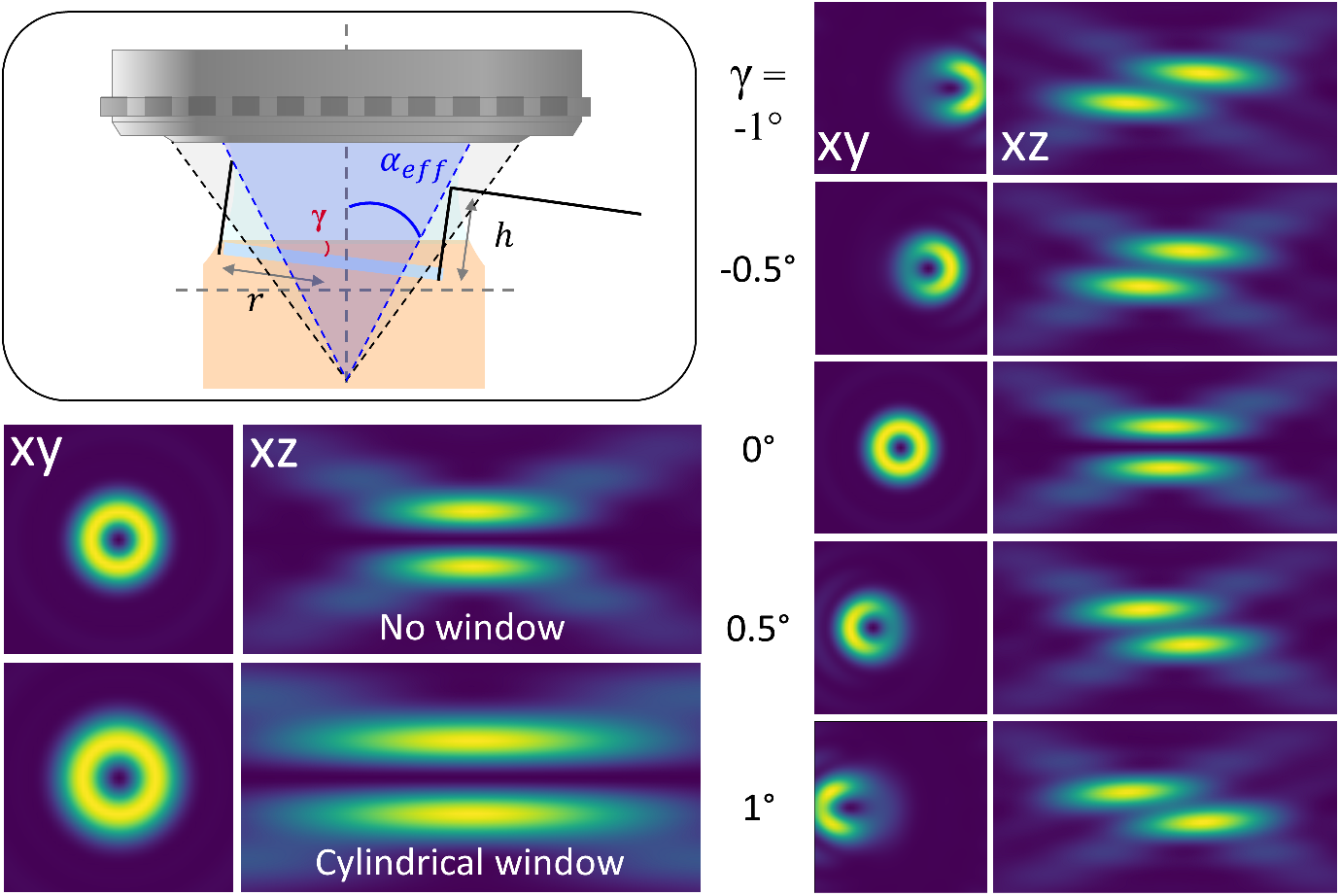
Focusing geometry. a) Schematic of the optical geometry after the microscope objective. b) PSF simulation with and without a cylindrical cranial window [23]. c) Impact of a tilted coverslip on the *donut-beam* profile. Image size 2 *×* 2 *µm*^2^ for XY views and 2 *×* 4 *µm*^2^ for XZ views.

Figure 4b illustrate the impact of a cylindrical cranial window. As described in [23], the presence of the window reduces the effective numerical aperture, leading to an extended focus. Additionally, a tilt angle of the coverslip introduces severe distortions of the PSF ([22]). This parameter is particularly critical *in vivo*, where a tilt angle ∼ 1^*◦*^ is inadvertently introduced.

#### 5.1.3 Input beam parameters

##### Intensity distribution, polarization and phase mask

The third tab in the main widget is used to define the properties of the input beam. The intensity profile incident on the input pupil of the objective is defined as a Gaussian profile with a specified wavelength and waist. Additionally, a misalignment of the input beam can be modeled through an offset of the intensity profile. A “Show intensity” push-button allows visualization of the input profile on the back pupil of the objective lens (see Fig 5a).

**Figure 5:**
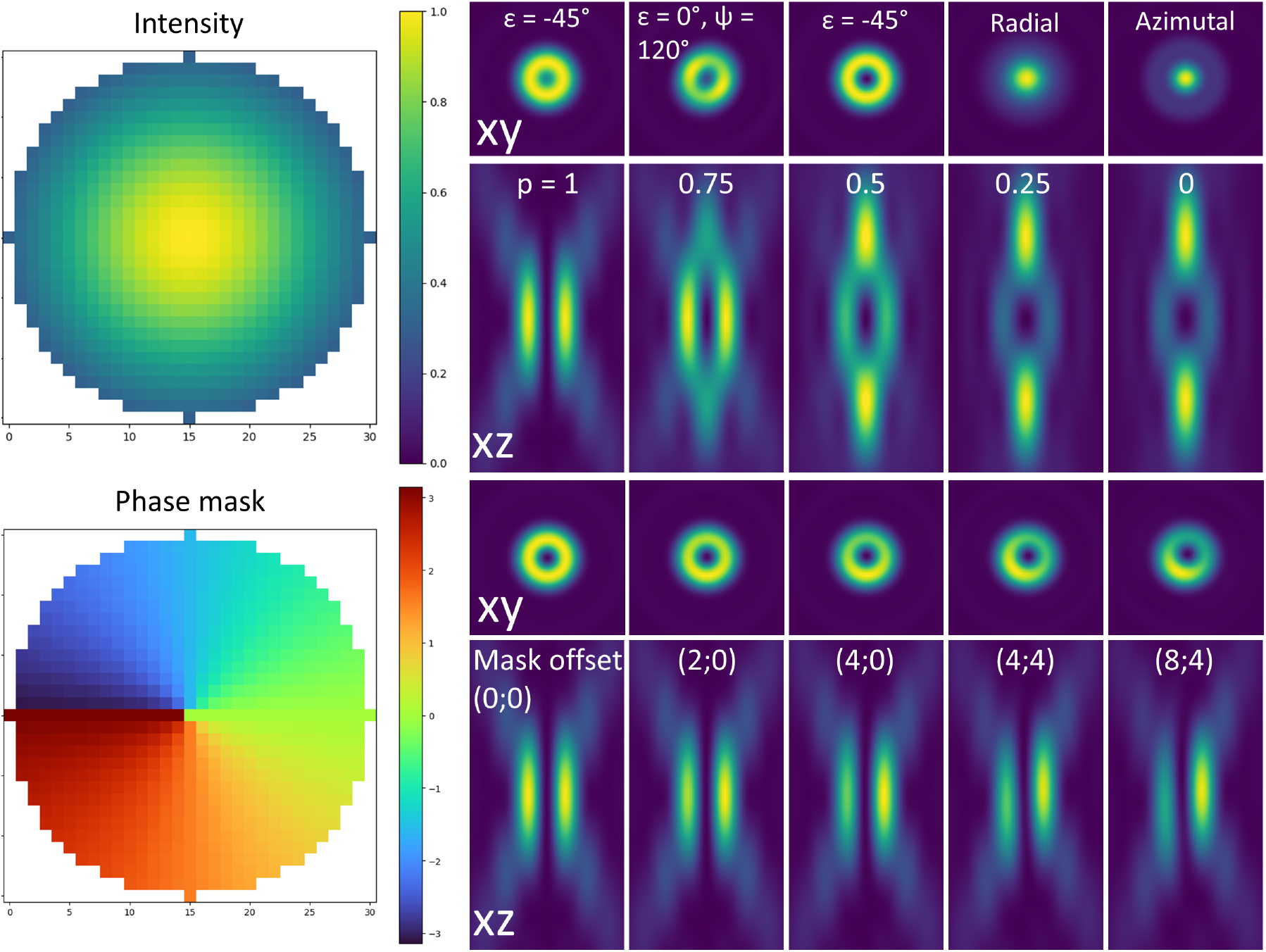
Input beam. a) Pop-up window displaying the intensity profile on the back pupil of the objective. b) Pop-up window showing the phase mask imprinted on the input beam. c) Effect of the polarization on donut beam PSF (in-plane views - 2 × 2 *µm*^2^). d) Donut and bottle profiles at different p ratios (axial views - 2 × 4 *µm*^2^. e) Impact of misaligned vortex mask on the donut-shaped PSF.

Faser also allows the user to set the polarization of the beam. A toggle box allows selection between elliptical (default), radial or azimuthal polarization. For elliptical polarization, two parameters (*ϵ*, Ψ) control the ellipticity (− 45^*◦*^ right-handed circular, 0^*◦*^ linear, and 45^*◦*^ left-handed circular).

Furthermore, a phase mask can be applied to the input beam. A toggle box allows selection of desired profiles (gaussian, donut, bottle or donut and bottle). Alternatively, a custom-defined phase mask can be loaded. Donut beams are formed using a vortex phase mask defined by an integer vortex charge. Bottle beams are associated with a ring phase mask defined by the ring radius and charge. In both cases, an offset can be introduced to model a misalignment of the phase mask with respect to the optical axis of the objective. Lastly, a combination of donut and bottle beams, commonly used in 3D-STED microscopy, can be modeled with a parameter p indicating the intensity split between the two phase modulations. Once again, a “Show Phase Mask” push-button enables to visualize the imprinted mask on the back pupil of the objective lens (see Fig 5b).

Figure 5 illustrates the impact of different parameters (polarization, phase mask offset) on the donut beam PSF. Noteworthy, as previously investigated [24], it clearly highlights that mask misalignment can easily be mistaken for optical aberrations.

##### Optical aberrations

Beyond the phase mask, to model a realistic optical system, optical aberrations can be added to the incident beam. We focused on the first 11 orders of the Zernike polynomial, as they represent the most important aberrations in optical microscopy. The fourth and final tab of the main widget allows independent setting of the coefficients for these 11 modes, as well as an offset of the aberrated wavefront. A push-button “Show aberrations” enables visualization of the input profile on the back pupil of the objective lens (see Fig 6a for an example).

**Figure 6:**
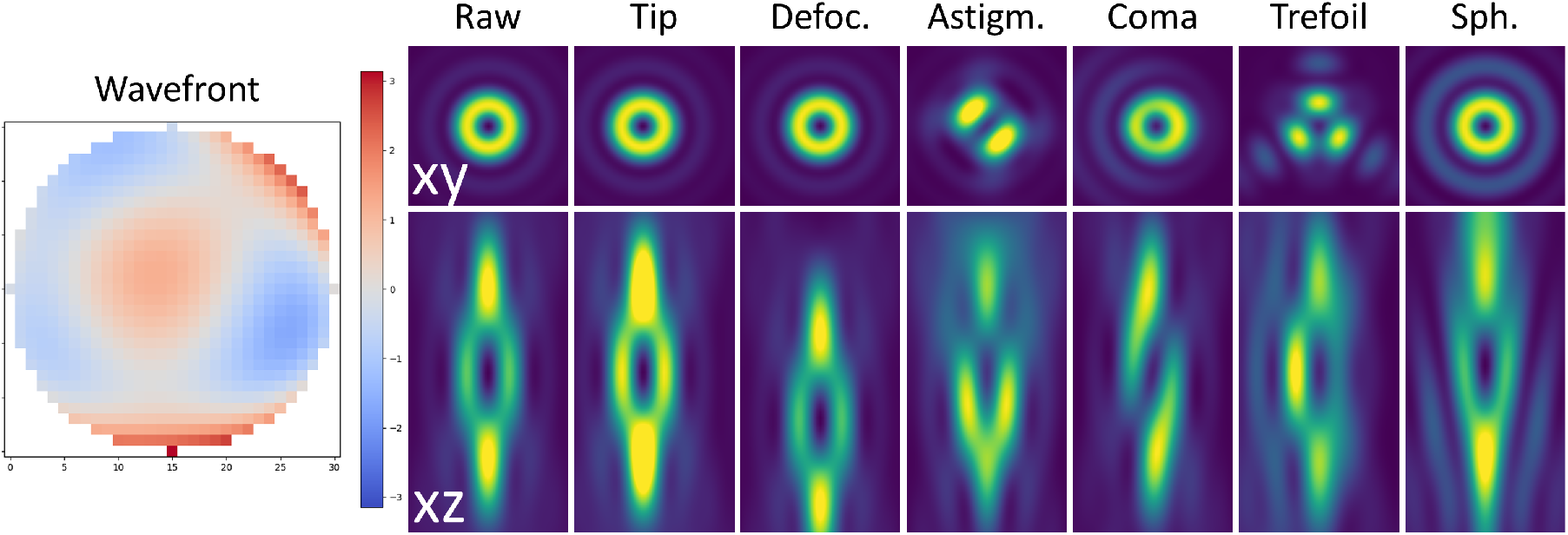
Optical aberrations. a) Pop-up window showing an example of an aberrated wavefront (a mixture of aberrations) on the back pupil of the objective. b) Impact of the main aberrations on donut and bottle PSFs (with *p* = 0.5). Lateral views 2 *×* 2 *µm*^2^ and axial views 2 *×* 4 *µm*^2^.

**Figure 7:**
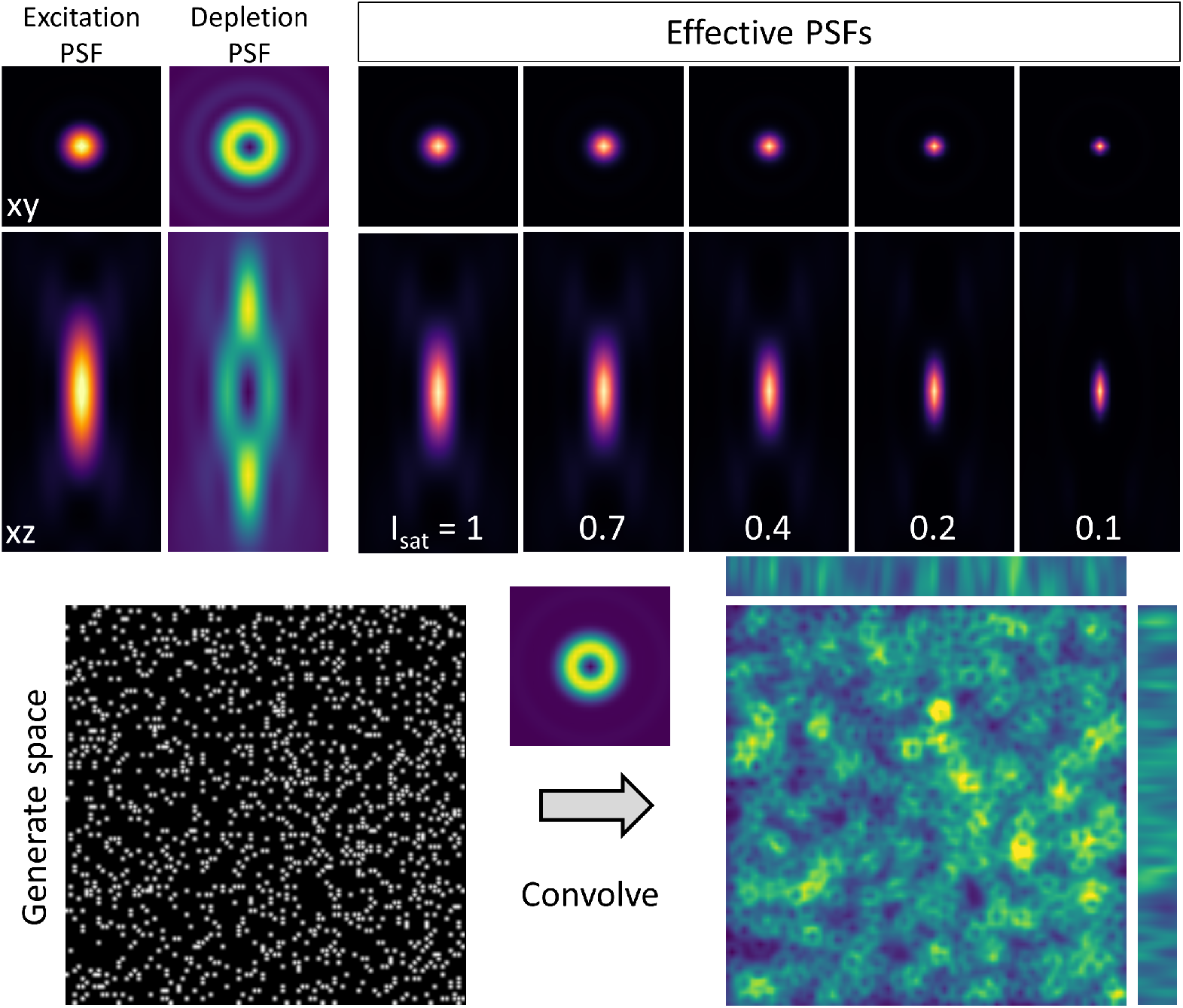
Helper widget. a) Example of an effective PSF calculated from the excitation and depletion PSFs (left panels) for various saturation factors. b) Example of a generated space (100 × 100 × 10 *µm*^2^) with 1000 spots. c) Convolution with the donut and bottle PSF.

Figure 6 illustrates the effect of the most classical aberrations on the mixture of donut and bottle beam profile.

### 5.2 Helper widgets

As explained above, the main widget serves to define simulations parameters for the PSFs. Faser also incorporates several helper widgets, providing additional functions.

A first tab enables to computes the effective PSF, in the case of STED microscopy. To do so, the user has to select exactly 2 previously simulated PSFs: one for excitation and one for depletion. The effective PSF is computed using the defined saturation factor according to eq. 24.

A sample tab enables to simulate an artificial sample of defined size in all three dimensions, with a specified number of points randomly distributed. The next tab serves to convolve images. It is originally intended to convolve a simulated PSF with a sample space, offering an illustrative view of the impact of a modified PSF on the image. Yet, it could also be used convolves two PSFs, notably to simulated excitation and emission PSFs,

Lastly, an export tab allows to save a z-stack of the simulated PSF as a .tiff file. This includes saving the simulation parameters as metadata accompanying the data.

## 6 Conclusion

In summary, we present Faser, a Python-based software package for simulating microscopy PSF using fully vectorial calculations. Faser is equipped with a comprehensive set of features that allow users to configure key optical parameters of the input beam (such as polarization and aberrations), as well as geometrical parameters related to the sample. In addition, it supports the simulation of wavefront shaping through the introduction of a phase mask. The software is intended as a pedagogical tool, enabling users to explore and better understand the impact of potential misalignments on microscope performance, with the goal to help identify the most important parameters that influence imaging quality.

## 7 Acknowledgements

This project received funding from the European Union Horizon 2020 research and innovation program under the *Marie Sklodowska-Curie Grant*, Agreement No. 794492, and from the *Fonds AXA pour la Recherche, AXA Banque Direction - Banque Patrimoniale et ses donateurs* to SB. Additional support was provided by the *Doctoral School for Health and Life Sciences* of the University of Bordeaux to JR, the *European Research Council* (ERC-SyG ENSEMBLE # 951294), *Human Frontiers Science Program* (# RGP0036/2020), *ERA-NET NEURON* (ANR-17-NEU3-0005), the *Féderation pour la recherche sur le cerveau (FRC)* and *Agence Nationale de la Recherche* (ANR-17-CE37-0011) to UVN.

## Notes

### Competing Interest Statement

The authors have declared no competing interest.

